# Modeling phytomeric growth using high-throughput phenotyping deconvolutes genetic control of complex traits underlying size and water use in *Setaria*

**DOI:** 10.1101/2023.12.28.573563

**Authors:** Renee Dale, Darshi Banan, Scott Oswald, Britney Millman, Haley Schuhl, Noah Fahlgren, Shankar Mukherji, Ivan Baxter

**Author notes:** Correspondence to (S.M.), (I.B.).

## Abstract

Plant growth and resilience is greater than the sum of its component traits, with important traits influencing one another. This interdependency makes it challenging to identify the genetic determinants of key agronomic traits, such as water use efficiency. Measuring traits such as plant height and size is possible via image analysis software, such as PlantCV, but these traits are often highly correlated. Furthermore, plant size is estimated using a 2D projection of a 3D object, which is more difficult for plants with complex body plans. To address these problems, we developed a generalizable, biologically-informed model describing the temporal coordination of semi-sequential phytomeric growth in the model grass *Setaria*. Our approach integrates time dependence with water usage, the growth of phytomers, and the emergence of side shoots or tillers, improving our ability to estimate both phytomer-level phenotypes and water usage traits in high-throughput. We developed new PlantCV methods to inform the Phytomeric Growth Model, and estimated parameter traits using a high-throughput phenotyping dataset of a recombinant inbred line (RIL) panel in well-watered and drought conditions. Model parameter estimates identified additional quantitative trait loci (QTL) for new traits compared to the directly measured PlantCV-derived estimates, and predicted the relationships underlying QTL control of composite traits through co-localization of model parameters to loci controlling size and water use metrics. The Phytomeric Growth Model estimates improved our ability to estimate the growth of tillers and obtain a model-based estimate of marginal water use efficiency (defined here as the ratio of biomass gained per gram of water transpired), both identified as highly influential on plant drought responses through random forest classification. Our approach demonstrates the value of integrating mechanistic modeling with high-throughput imaging to extract new information at scale.

## Introduction

Phenotypic variation emerges from interactions between the genome and the environment, but connecting the phenome to the genome is a major challenge in plant biology (Pauli et al. 2016). High-throughput phenotyping (HTP) is often implemented as a way to non-invasively capture developmental progression and phenotypic variation in large populations (Danilevicz et al. 2021; R. Wang et al. 2020; Ferguson et al. 2021; Varela et al. 2022). This approach enables the phenotyping of hundreds of genotypes in a single experiment, generating the statistical power required to identify genetic loci driving traits of interest.

However, HTP approaches are limited in what traits they can measure, and measurable traits can be difficult to connect to specific plant processes. For example, computer vision directly and successfully measures overall area, and height (R. Wang et al. 2020; Gehan et al. 2017), and robotic weighing and watering systems can track biomass production and how much water is being used by each plant. Yet these traits emerge from diverse plant processes across scales and time and none of them have been completely described at the genetic level. Understanding the more proximal processes driving changes in downstream traits is important for improving plant performance.

One under-studied contribution to plant performance is the impact of developmental transitions.Developmental transitions are not typically integrated into models that address key agronomic traits. However, we know that developmental transitions impinge on important traits. Proximal processes involved in developmental progression in grasses like *Setaria*, such as leaf elongation and phytomer emergence rates, can explain differences in biomass allocation, size, and height between genotypes or across environmental conditions (Voorend et al. 2014; Zhang et al. 2016). Leaf elongation rate has been used extensively to characterize the growth of a phytomer (Xue et al. 2012; Fournier 2000; Fournier et al. 2005; Zhang et al. 2016; Reddy et al. 1997) and along with leaf emergence rate (the rate at which nodes are produced), is a major determinant of leaf area. The temporal coordination of developmental progression is highly controlled and responsive to environmental stressors (Hodge and Doust 2021; Fournier 2000; Fournier et al. 2005; Litvin et al. 2016; Coussement et al. 2021; Sasidharan et al. 2025) and understanding how developmental transitions are coordinated, along with the allocation of biomass to individual organs, are current challenges in plant developmental biology and physiology (Poethig 2003; Poethig and Fouracre 2024; Zhang et al. 2016; White et al. 2016).

Despite the importance of describing the temporal coordination of developmental processes, phytomeric growth rates have largely not been represented in computational models that can help quantify novel traits that capture key information on plant allometry and growth. Often, the semi-sequential emergence and growth of individual phytomers are modeled as pooled composites (Matthews et al. 2021), or as discrete events (Fournier 2000; Xue et al. 2012). Ideally, developmental processes and WUE could be integrated in a biologically-inspired manner with mechanistic systems modeling.This approach is thought to lead to better understanding of the processes underlying allometry (Hodge and Doust 2021) and is a more appropriate structure than assuming properties of individual growth rates (Coen and Prusinkiewicz 2024). Well-specified systems models can contextualize data emerging from the non-linear feedback loops often observed in biological systems (Lucido et al. 2025; Monsalve-Bravo et al. 2022). More broadly in biology, multi-scale and mechanistic models are able to contextualize complex trait relationships during development (Danilevicz et al. 2021; Benes et al. 2020; Coussement et al. 2021), but are underutilized in plants (Lucido et al. 2025; Dale et al. 2021).

One critical trait influencing plant performance is water use efficiency (WUE), the ratio of the rate of carbon gain to the rate of water use (Liang et al. 2023; Bhaskara et al. 2022). However, quantitative descriptions of WUE can be challenging to produce for a number of reasons (Liang et al. 2023; Leakey et al. 2019). Photosynthetic CO_2_ assimilation and transpiration are highly sensitive to environmental fluctuations, and, while leaf-level WUE is readily assessed at short timescales, whole-plant WUE over the life of the plant has traditionally been more challenging to quantify (Liang et al. 2023). While whole-plant WUE can now be estimated in HTP systems by dividing the estimated plant size (e.g., pixel area) or biomass by the total amount of water used, plant size and water use are highly correlated, limiting the observable variation in WUE (Halperin et al. 2017; Feldman et al. 2018). This hampers our ability to identify novel genetic loci that independently contribute to genotypic differences in WUE or to plant size. Finally, where imaging is used rather than lysimetry, pixel area is a 2D projection of a 3D object that includes tissues of different densities, further complicating the accuracy of biomass and productivity estimates (see Fig. 1A).

**Fig. 1.**
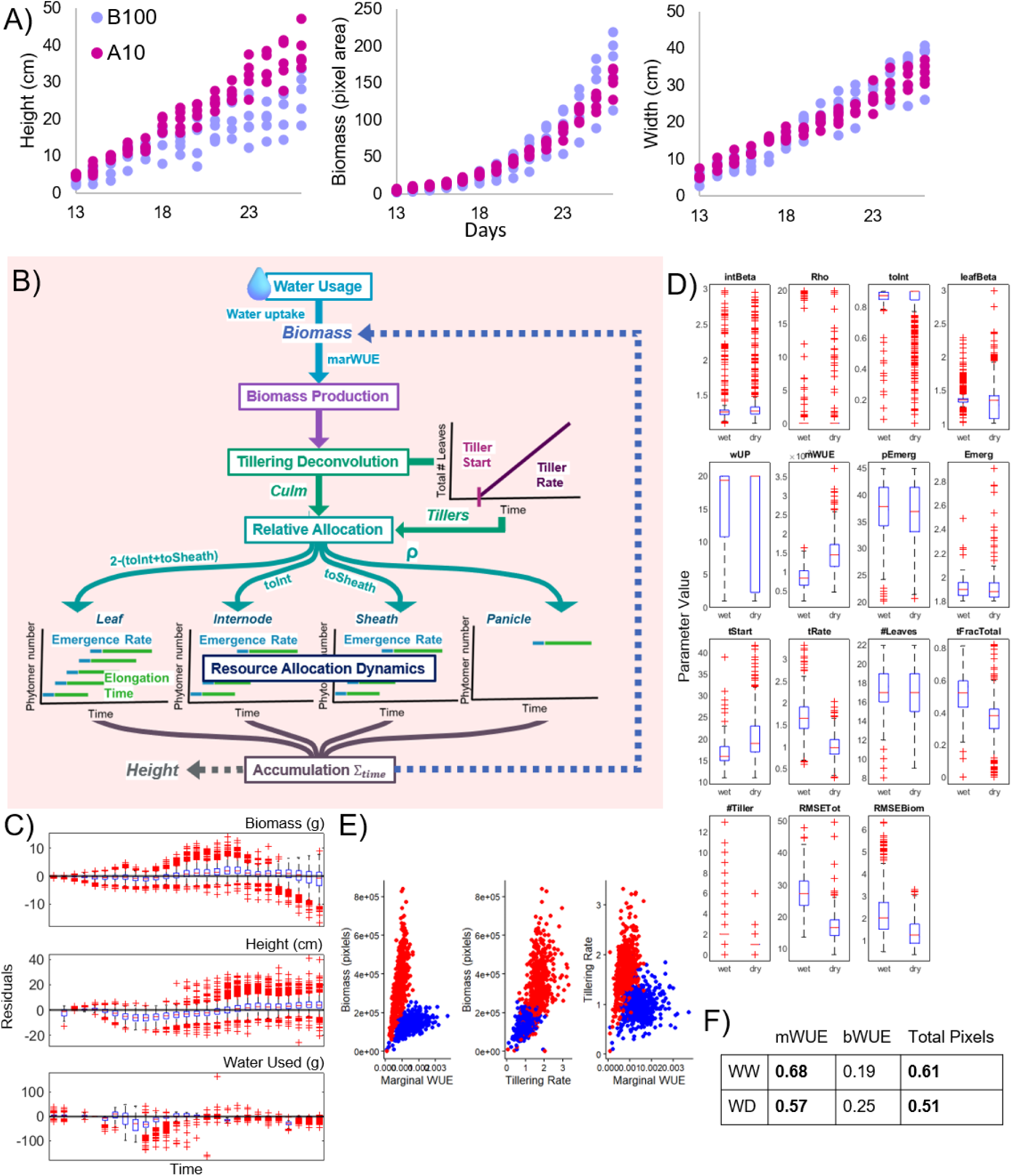
Developmentally-informed mathematical Phytomeric Growth Model enables high-quality estimates of phenotypic traits. A) Imaging-based estimates of plant phenotypes can be misleading. Time series of the growth of 5 wild A10 and 5 domesticated B100 *Setaria* from the HTP platform are shown for plant height, biomass, and width. Although A10 is very bushy with lots of tillers, while B100 is a domesticated, both genotypes appear similar. B) A mathematical model of *Setaria* growth based on the temporal coordination of the semi-sequential growth of phytomers. The model consists of seven modules (visualized as boxes). Available water and current leaf mass (culm or tiller) determine biomass production over time (*Water Usage* module). Tiller emergence is described by the *Tillering Deconvolution* module. Biomass produced is allocated to phytomer components (leaf, internode, sheath, or panicle) by the *Relative Allocation Fraction* module. The *Resource Allocation Dynamics* module determines the number and type of phytomer component growing at each time point, determined by the overall phytomer emergence rate and the elongation duration parameters. Finally, the *Allocation* module calculates the resources allocated over time to form composite traits such as leaf biomass, an input to the *Water Usage* module, making the model closed-loop and self-contained. Boxes indicate model modules. Italics indicate biological components in the model. Parameter values describing modules are shown adjacent to the relevant module. Solid lines indicate flow of information, and dashed lines represent composite calculated traits. Model description is provided in Supplemental File 1. C) Phytomeric Growth Model fit residuals as a function of time. Residuals (calculated here as the difference between the model and high-throughput phenotyping data) obtained during fitting the model to the HTP data (Feldman et al. 2017) are plotted. Top to bottom, panels show residuals in biomass (g), height (pixels), and water used (g). Zero indicates perfect agreement between model and data. Representative model fits to four random plants in the HTP dataset are provided in Supplemental Figure 4. Blue boxes indicate 25th and 75th percentiles. Red line indicates the mean, 50th percentile value. Black whiskers indicate 1 times the inter-quartile range. Red crosses indicate outliers beyond the whiskers. D) Parameter estimates of phenotypic traits in well-watered and drought conditions across all genotypes. Model parameter estimates resulting from fitting to HTP data (1C, (Feldman et al. 2017) are shown as box plots across genotypes for the two treatment conditions. Blue boxes indicate 25th and 75th percentiles. Red line indicates the mean, 50th percentile value. Black whiskers indicate 1 times the inter-quartile range. Red crosses indicate outliers beyond the whiskers. E) Model-based estimate of marginal WUE, tillering rate, and size cluster by treatment. Model-estimated marginal WUE and tillering rate were identified as influential parameters describing plant size across all genotypes and treatment conditions. Influential parameters are identified via random forest, see Methods and Supplemental Fig 6-7. Red indicates drought, and blue indicates well-watered conditions. F) Correlation coefficients illustrate estimate quality. The Phytomeric Growth Model-estimated marginal WUE (“mWUE”) has a stronger correlation coefficient with the ground truth (dry weight) than both the whole-plant WUE estimate (“bWUE”) and biomass estimated by pixel area. The bWUE is estimated by dividing total pixels by total water (g) provided during the course of the experiment. Moderate (0.5 - 0.7) correlation coefficients are bolded. Correlations are separated by well-watered (WW) and drought (WD) conditions.

Here, we introduce a mathematical model of the temporal coordination across phytomeric development in *Setaria* that integrates the usage of water (e.g., water uptake rate, water use efficiency) with developmental progression (e.g., phytomer emergence, tillering). We found that our Phytomeric Growth Model parameter traits improved correlations to ground-truth plant size, improved our ability to map alleles underlying genotype to phenotype relationships compared to the initial computer vision trait estimates, predicted phytomeric links underlying QTL control of composite traits, and found new QTL controlling traits estimated by model parameters.

## Results

### Mechanistic model development

In order to measure new developmentally-informed phenotypes, we developed a Phytomeric Growth Model to describe the temporal coordination of semi-sequential phytomeric growth exhibited in *Setaria* as a case study for other grasses (e.g., maize (Fournier 2000)). *Setaria,* a well studied model C4 grass (Feldman et al. 2017, 2018; Doust et al. 2019; Hodge and Doust 2021; Brutnell et al. 2010), can produce several shoots: the main shoot (or culm), and additional shoots (i.e., tillers) which emerge later.

The Phytomeric Growth Model consists of seven modules describing separate processes (Fig. 1B). The model’s core module, the *Resource Allocation Dynamics* (*RAD*, Fig. 1) defines a simple set of rules for *Setaria* phytomer development. Under semi-sequential growth, a shoot produces a new phytomer at the phytomer emergence rate (**τ**), and each phytomer reaches maturity after a certain elongation duration (**β**, which is proportional to the plant’s age when the phytomer is produced, **τ**\***β**). These parameters determine when a phytomer is growing and how long its components (leaf blade, sheath, internodes, and panicle) grow, and thus how many phytomers are growing concurrently, including tiller phytomers. These parameters alone do not determine the final size of the phytomer (see Supplemental Text 1), as it is influenced by water use and the growth of other phytomers. The production of tillers is described in the *Tillering Deconvolution* module (see below).

In addition to the number of growing phytomers, the *RAD* module uses the *Relative Allocation Fractions* (*RAF*) module to determine how much of a given resource is allocated to each phytomer and to each of its components at each time point, weighted by the number of growing phytomers (see Supplemental Text 1 for details on model formulation). Additional growing phytomers increase the competition for resources among those phytomers, lowering the relative allocation to each phytomer. Completed phytomers supply resources to growing phytomers. Thus, variation in the elongation rate of phytomer components emerges as a result of the interaction between resource production dynamics and developmental progression dynamics.

The *Water Usage* (*WU,* Fig. 1B) module determines the rate of biomass production at each time point using a system of five ODEs. This sub-model takes as inputs the water provided on each day and the current RAD-derived estimates of tiller and culm leaf mass. The outputs of the *WU* module are biomass production and water usage for each day. Biomass production is fed into the *Tiller Deconvolution* (*TD*) module, which estimates the number of tillers produced via a linear model (see Supplemental File 1 for more details on model formulation). The current estimated number of tillers are then provided to the *Resource Allocation Dynamics (RAD)* module, which determines what fraction of biomass production is added to each phytomer and tiller. The resulting leaf biomass is provided to the *WU* module to compute the next day’s production.

Leaf biomass is used as the imaging process cannot estimate leaf biomass and leaf area independently. A subset of plants were harvested to measure leaf mass and total biomass to determine the relationship between observed green pixel area (a proxy for leaf area) and leaf biomass (Feldman et al. 2018). Water usage is calculated per-leaf mass (in grams):

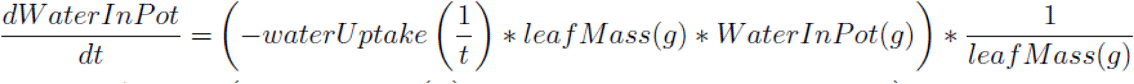

The amount of water in the pot is reduced at a rate of waterUptake, and calculated per leaf mass. The initial conditions (total amount on each day) of WaterInPot on a given day D are *WaterProvided (D) + WaterInPot(D-1)*. The amount of water used on a given day D is calculated as *WaterInPot(D) - WaterInPot(D-1).* This value is compared to the data on water usage over time.

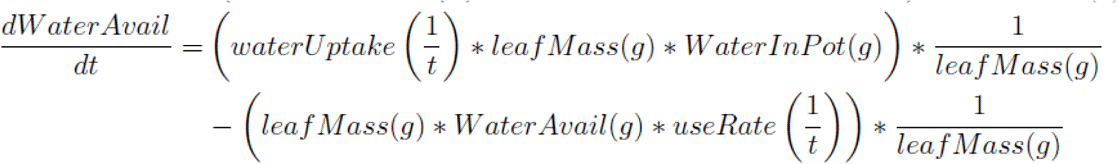

Water is made available to the plant at the rate of waterUptake and modulated by the available WaterInPot. This is calculated per leaf mass. Water is used at a useRate modulated by the total waterAvailable, also calculated per-leaf mass. We assume that all water available to the plant is used at each time point (*useRate=1*), an assumption that works well when plants are sufficiently large to access the entire volume of the pot. The model assumes that all water lost from the pot is transpired via the leaves—-soil evaporation accounts for about 10% of water loss from the pot, but this is assumed to be the same for all pots and implicitly accounted for in the water uptake rate parameter.

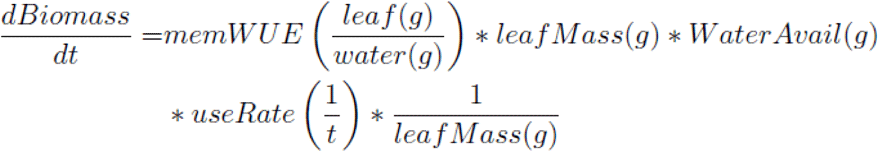

Biomass produced at each time point is calculated per-leaf mass, which is determined as

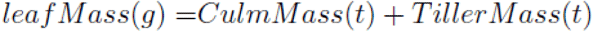

The initial CulmMass and TillerMass at the beginning of each daily simulation (when new water is added) is determined by the culm leaf and tiller leaf mass estimated by the *RAD* module. Inter-simulation culm and tiller leaf mass is approximated by including all accumulated mass. Biomass production is calculated as the water in-flux (waterAvail *useRate, per leaf mass) multiplied by the model-estimated marginal water use efficiency (memWUE), giving us grams of biomass.

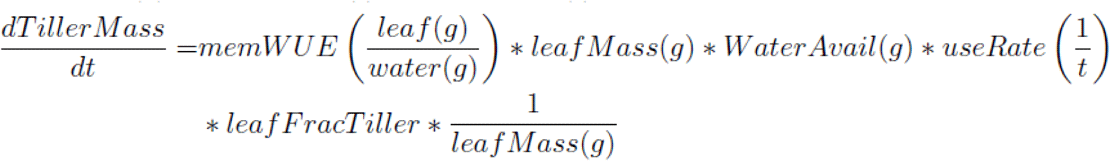

The amount of tiller mass produced per time point is calculated as the biomass production at each time point multiplied by the leafFracTiller parameter. LeafFracTiller is calculated based on the current total plant shoots (culm and any tiller shoots). We found that biomass must be shared by number of shoots, rather than relative biomass, for newly emerged tillers to grow while the culm is relatively much larger. While the culm is growing,

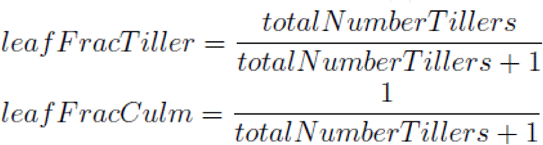

Culm mass production is calculated in the same manner.

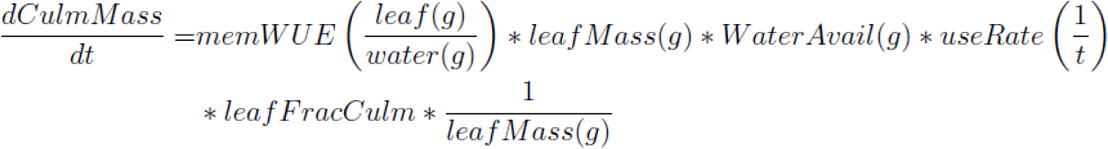

The parameters memWUE and waterUptakeRate are estimated during the fitting process along with the remaining parameters describing the other modules. The ODE-based calculations of CulmMass and TillerMass gains are provided to the RAD module, which in turn estimates the amount of biomass (in grams) allocated to tiller leaves and culm leaves. The summation of these is compared against the Bellwether imaging estimates of plant biomass. The *RAD* module estimates of stem height are compared to the Bellwether imaging estimates of plant height.

As *Setaria* produces tillers which compete for the same resources as the growing phytomers on the culm, the *Tillering Deconvolution (TD,* Fig. 1B) module calculates the number and size of tillers using the tiller start time parameter (the plant’s age at which it produces its first tiller) and the tillering rate parameter (the rate at which new leaves are produced). New tillers emerge as the number of predicted emerged leaves over time exceeds the number of leaves in existing culm and tiller(s) (predicted using **τ** and panicle emergence parameters). Each tiller is modeled using the same developmental parameters in the *RAD* as the culm, but temporally delayed (Mauro-Herrera and Doust 2016). The amount of resources allocated over time are accumulated to give predictions of phytomer lengths, which are integrated to provide composite trait predictions like height (here, internodes and sheaths), and leaf biomass. Total leaf biomass determines water usage and biomass production, making the model fully self-contained and a closed-loop. The model takes water provided data as an input, and calculates water usage, biomass production, growth, and allocation to each phytomer (leaf, sheath, internode, and panicle) for each day in the simulation. These calculated values are allowed to vary within bounds, and are uniquely estimated for each plant sample. The model predicted total biomass (sum of leaf mass) and height (sum of internodes and sheathes) are compared against the data during parameter estimation.

The resulting Phytomeric Growth Model provided a novel approach to describing the temporal coordination of growth across phytomers using coupled developmental rates, biomass production and allocation. To test the appropriateness of the *RAD* and *RAF* modules, and obtain initial estimates of the developmental parameters, we performed a validation experiment. Fifteen domesticated (accession B100) and fifteen wild (accession A10) *Setaria* plants were grown in the greenhouse and the required model parameters were measured by hand (see Methods). We tested the model’s prediction of resource allocation among phytomers by comparing the relative length of each phytomer in simulations to the observed relative distribution, and found good agreement between the two. The data collected also gave us insight into the novel parameter traits, the phytomer emergence rate, and the grow time durations of leaves and internodes (See Supplemental Figures 1 and 2).

**Fig. 2.**
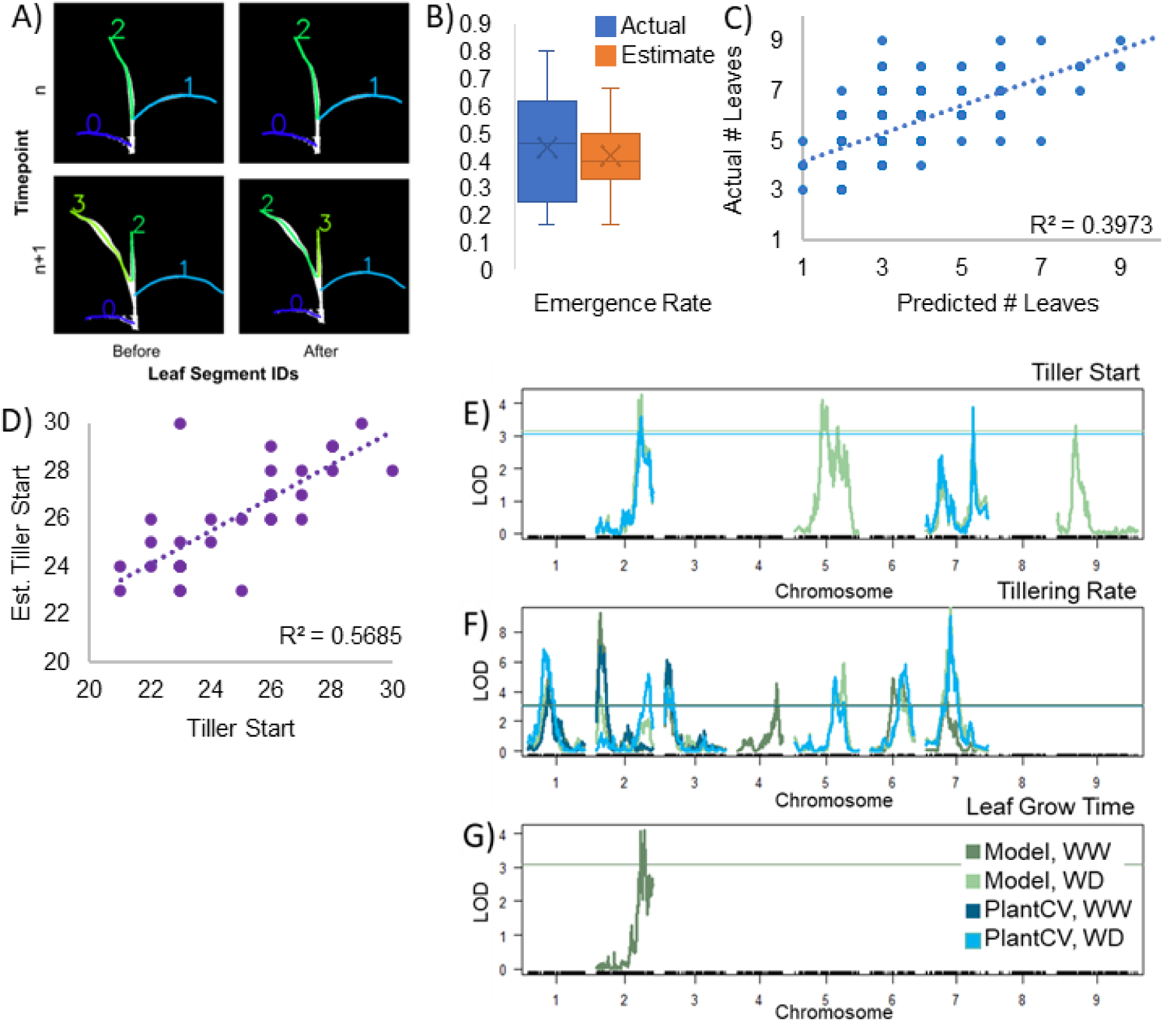
PlantCV provides initial estimates further refined by the mechanistic Phytomeric Growth Model. (A) Leaf identities in PlantCV before and after the novel implementation of leaf branch point location sorting. The numerical identity of leaves on each imaging sample were tracked over time, using node locations (the branch point of each segment identified during segmentation) to inform identity. This allowed the estimation of traits such as leaf emergence rate and leaf elongation duration using the individual leaf lengths over time, until the start of tillering. B) Plant-CV derived estimates of the phytomer emergence rate (here, as days per phytomer) are consistent with the manual measurements on average. C) There is a moderate correlation between the actual and predicted number of leaves. D) There is a good correlation between estimated and actual tiller start time. (E-G) Phytomeric Growth Model refinement of trait estimates provide additional QTLs in tiller start time (B), tillering rate (C), and leaf grow time duration (D). The model provides us with two additional QTL controlling tillering intuition, one additional QTL controlling tillering rate, and three additional QTL controlling leaf grow time. PlantCV estimates of tillering rate provide two additional QTL. Model-derived parameter estimates are shown in green, PlantCV-derived traits in blue. Traces in darker colors indicate well-watered conditions. PlantCV-derived traits are used as initial guesses that the model then refines using all available data in the global optimization procedure. No QTL were found for the other PlantCV-derived trait, the phytomer emergence rate.

### The model estimates water usage traits in the context of phytomeric growth

Increased growth increases total transpiration, producing a strong correlation between water use and biomass in both well-watered and drought conditions (Feldman et al. 2018). As a result, estimating whole-plant WUE as the size divided by the water used produces another strongly correlated metric that limits our ability to identify novel genomic loci. Our Phytomeric Growth Model extracts additional biological insight by quantifying the interrelatedness between plant traits. The model describes the temporal coordination in phytomeric growth, with the *Water Usage* module describing soil water uptake and biomass production (Figure 1B). Since our model-derived metrics of water use are estimated in the larger context of a mechanistic multi-scale model (Figure 1D), we provide evidence for a relationship between tillering and water use efficiency (Figure 1E), and our model-based marginal WUE estimate correlates more strongly with the dry weight compared to an imaging-derived estimate (Figure 1F, Supplemental Table 5). The marginal WUE calculated by our model describes the tradeoff between water loss and carbon gain in the context of a whole system incorporating biomass production, phytomeric growth, and tiller emergence.

### Informing the Phytomeric Growth Model with PlantCV

The *RAD* module at the core of our mathematical model calculates the final leaf size using the phytomer emergence rate and the elongation duration of each phytomer component (leaf blade, sheath, and internode) in the context of water uptake and growth. While these parameters can be measured by hand, the time required makes measuring a full set of over 1000 plants unfeasible (at two minutes per plant per timepoint, hand-measuring would require around 40 person hours every timepoint). PlantCV is an image analysis software package used to extract trait estimates in high throughput. To fully inform the Phytomeric Growth Model, current PlantCV capabilities were expanded to obtain as many estimates for the model parameters as possible by implementing leaf identity tracking via branch point (collar) location sorting (Figure 2A). This provided a proxy for phytomer emergence over time (Figure 2B), and enabled estimation of the leaf elongation duration by tracking the number of days from phytomer emergence until a given leaf reaches terminal length. This also provided us with total leaf number estimates (Figure 2C).

Using computer vision to automatically identify when each leaf emerges on the culm is complicated by tillering. Newly emerged leaves of tillers may be mistaken for newly emerged culm leaves. The tiller start time (Figure 2D) was estimated using an existing PlantCV function to count segment ends. Tiller start time is defined as the time when two or more leaves were detected as emerging in a single day, as our validation experiment suggested a rate of one leaf emerging every two days. The phytomer emergence rate and leaf elongation duration are then constrained to be estimated using data prior to the date of tillering initiation. The tillering rate is obtained using a linear regression to the number of leaf tips over time. A slope of ∼1 corresponds to 0-1 tillers, which is domesticated B100-like; and a slope of ∼2 corresponds to many tillers, or wild A10-like (see Figure 1D, Supplemental Table 3 for across-genotype information). The slope and intercept of this regression is used in the mathematical model in the *Tillering Deconvolution (TD)* module. Using these estimates, the *TD* module determines the initiation of additional tillers by calculating the emergence of new leaves.

### Biological insight using parameter traits

Previously, a study on 1138 plants from 185 genotypes in a *Setarial RIL* population under well-watered and drought conditions identified plant size, height, and three water use metrics co-localizing onto four QTL (Feldman et al. 2018, 2017). To obtain biological insight on using the Phytomeric Growth Model, we obtained parameter trait estimates on that dataset. Global optimization via Matlab MultiStart was used to estimate parameter traits on each sample (biomass, water use, height) across 200 starting points that included PlantCV-informed initial guesses when available (see *Methods: Parameter trait estimation* for more detail. The full set of parameters and descriptions is included as Supplemental Table 1). This provides plant and genotype-level estimates of parameter traits, including *RAD* traits (phytomer emergence, leaf grow time, sheath grow time, internode grow time, panicle emergence time); *RAF* traits (toInt, toSheath, rho), *TD* traits (tiller start time, tillering rate), *WU* traits (water uptake, marginal WUE); and model-derived calculations (number of leaves, tiller fraction of total mass, and number of tillers). Descriptions of model parameter estimated values are in Figure 1C, Supplemental Tables 1 and 4.

### Phytomeric Growth Model improves the ability to map on to new PlantCV traits

Four parameter traits were estimated using PlantCV: the phytomer emergence rate, leaf elongation duration, tillering initiation time, and tillering rate. We compared the QTL mapping between the raw PlantCV estimates (missing and unrealistic estimates removed, see Methods and Supplemental File 3) and the Phytomeric Growth Model-derived estimates, which use PlantCV estimates as initial guesses during the global optimization process. Most of the QTL were significant using either estimate (Figure 2E-G). The Phytomeric Growth Model-based estimates of the tiller start time provided two additional QTL (5@59, 9@39). The model-based estimates of the tillering rate provided three unique QTL (4@81, 5@100, 6@48) compared to one unique QTL (2@111) by PlantCV estimates, with 5 loci identified by both estimates. The model-based estimates of the leaf grow time produced one QTL not found using PlantCV estimates (2@101). The leaf grow time trait was difficult to estimate using PlantCV, as the time when 2-3 leaves reached terminal length often coincided with tillering and leaf overlap. No QTL controlling the phytomer emergence rate was found using either approach, likely because this value was highly consistent across genotypes and experiments in our conditions (Supplemental Fig 1A-B).

### The Phytomeric Growth Model partitions the QTL controlling size and water use into more specific phytomeric parameter traits

Phytomeric parameter traits influencing size (e.g., leaf grow time, tillering rate), water usage (water uptake, model-estimated marginal WUE), and height (internode allocation fraction) co-map to the four QTL in different combinations (Figure 3A). The Phytomeric Growth Model predicts the mechanisms underlying the co-localized control through the phytomeric traits that are also controlled by the four QTL. The QTL at chromosome 2 and at position 96 centimorgans (2@96) was found to control the model estimate of marginal WUE, leaf elongation duration, and tiller start time. The QTL at 5@100 was found to control the water uptake rate, the model estimate of marginal WUE, tillering rate, tiller fraction of total mass, and number of tillers. The QTL at 7@100 was found to control the internode allocation fraction, tiller start time, and tiller fraction of total mass. The QTL at 9@35 was found to control the water uptake rate, the model estimate of marginal WUE, tiller start time, and tiller fraction of total mass. Only the QTL on chromosomes 2, 5, 7, and 9 are shown here. Full QTL mapping results provided in Supplemental Table 2, and visualized in the context of the model diagram in Figure 3B.

**Fig. 3.**
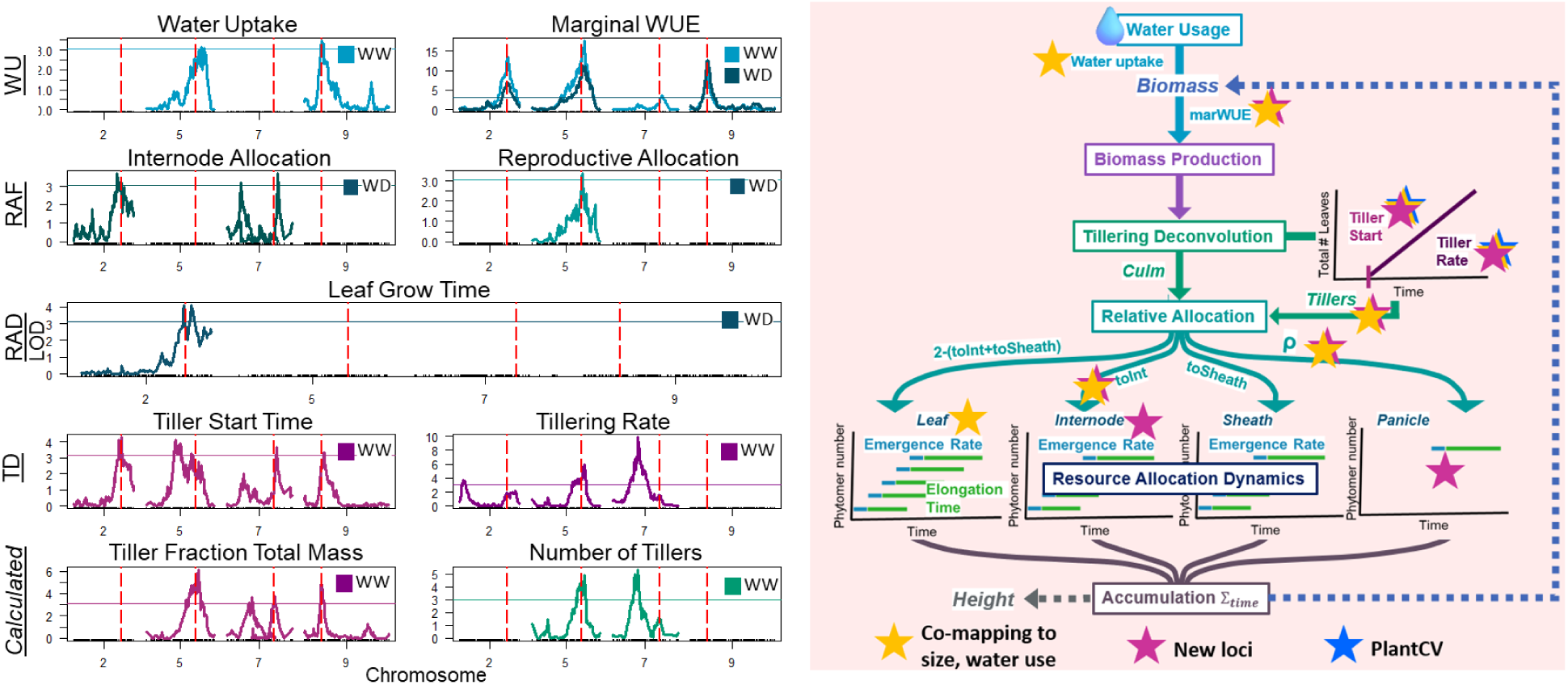
Parameter traits predict mechanisms underlying QTL control of composite traits. (Left) Previously four QTL (2@96, 5@100, 7@100, 9@35) were found to control both plant size and three metrics of water use (Feldman et al. 2018). The loci 5@100 was also found to control height in the same dataset (Feldman et al. 2017). The Phytomeric Growth Model predicts the mechanisms underlying this control through the phytomeric traits that are also controlled by the four QTL. Only four chromosomes are shown, and red dashed vertical lines indicate the location of the four QTL. Marginal WUE refers to our model-based estimate of mWUE. (Right) Contextualizing QTL control of parameter traits in the Phytomeric Growth Model. Gold stars indicate model parameter traits that correspond to the four QTL controlling composite traits (Feldman et al. 2018, 2017). Blue stars indicate traits estimated using only PlantCV that are controlled by new QTL. Pink stars indicate parameter traits with new QTL.

In addition to the four QTL found in Feldman et al., the parameter traits enabled the identification of 14 new QTLs (Supplemental Table 2). All parameters but one mapped to at least one QTL. The strongest of the new QTLs was for the tillering rate on chromosomes 7 (7@50), chromosome 2 (2@9), and chromosome 3 (3@5). The strongest of the new QTLs for non-tillering traits was the internode grow time on chromosome 5 (5@80).

We also obtained predictions on the influence of drought on phytomeric traits. Most model-based estimates of phytomeric traits changed significantly due to drought (Figure 1D), including an increase in the internode grow time duration, a decrease in the fractional allocation to the internodes, an overall reduction in tillering (increased tiller start time, reduced tillering rate, and a corresponding reduction in the tiller fraction of total mass and the number of tillers) (Supplemental Table 4 shows t-test results). The Phytomeric Growth Model predicts an increase in water uptake rate and in the model estimated marginal WUE under drought conditions. Details on fitted model parameter trait values are provided in Supplemental Table 3.

## Discussion

In this paper we develop a novel mathematical model describing the temporal coordination in semi-sequential phytomeric growth in *Setaria*, enabling the estimation of critical traits like the marginal WUE in context of the complex relationships and feedbacks. This Phytomeric Growth Model consists of four core assumptions: that the plant produces a new phytomer at a fixed rate, that each phytomer grows for a set duration independent of biomass production, that the final size of each phytomer depends on the overall productivity of the plant during the phytomer’s period of maturation, and that growing phytomers compete equally for photosynthate. These assumptions enable the coupling of overall growth to developmental progression. Core model parameters include developmental time parameters, such as the phytomer emergence rate and elongation duration, which are major determinants of plant size (Voorend et al. 2014). We hypothesized that this approach would improve our ability to estimate critical traits like marginal water use efficiency through mechanistic representation of core growth processes that are not fully observable in the high-throughput system. While water use efficiency can be estimated at the whole-plant level by taking the ratio of the green pixel area and the total water used during the course of the experiment, biomass estimated by green pixel area is a 2-dimensional projection of a 3-dimensional object, and size will be under-estimated, particularly in genotypes with higher tillering rates, and density differences between leaves and stem are ignored. The mechanistic structure of our phytomeric Phytomeric Growth Model enables the capture of the progression of development in both the culm and any tillers.

We tested our hypotheses using an existing dataset (Feldman et al. 2017, 2018), where four QTL were previously found to control composite traits (plant size and four water use metrics) on a *Setaria* recombinant inbred line (RIL) population. We analyzed these data using our Phytomeric Growth Model and performed a QTL analysis to determine 1) how modeling water use and the progression of development would influence trait estimate quality; 2) how model-based parameter trait estimates mapped to QTLs would compare to computer vision-based estimates; 3) what parameter traits would map to the four QTL controlling composite plant phenotypes (height, size, water use); and 4) what, if any, parameter traits would map to new QTL.

We identified the tillering rate and the model estimated marginal WUE as highly influential parameters driving plant response to drought. All wild grasses produce tillers, which are offshoots of the main culm. These offshoots produce roots that can eventually allow for growth independent of the culm (Mauro-Herrera and Doust 2016). Tillering drives increased plant size, and tillering rate influences drought resistance; low-tillering genotypes tend to have higher drought resistance (Tardieu et al. 2020; Alam et al. 2014), while higher tillering is more advantageous when water is available (Tardieu et al. 2020). Higher tillering is also associated with higher leaf area index (LAI), possibly contributing to a reduction in soil evaporation that improves drought resilience (Vadez et al. 2024).

Tillering-related phenotypes were well-represented in our mapped QTL. The Phytomeric Growth Model describes coupled biological processes that occur during development and growth, capturing constraints and feedbacks (e.g., tillering, leaf emergence, reproductive growth). In conjunction with PlantCV measurements, our model can distinguish a ‘large plant with no tillers’ from a ‘small plant with many tillers’ even when the overall biomass area and height of each appears similar. The Phytomeric Growth Model accounts for the effect of tillering on the water utilization of the culm, a model-based estimate of the marginal WUE, and overall plant size by estimating these factors in conjunction with each other in a biologically-inspired model structure. Four loci co-localized to plant size and water use traits (2@96, 5@100, 7@100, and 9@35) also co-mapped to tillering traits (Supplemental Table 2). Eight additional QTL were found to control tillering traits. Some of these QTLs overlap (+/- 10 centimorgans) with those found previously in another study on the *Setaria* RIL population (Mauro-Herrera and Doust 2016), specifically at loci 1@35, 2@9, 5@58, 5@100, 6@42, 7@100, and 9@36 for tillering-related phenotypes, and loci 2@86, 7@30, and 7@45 for height-related phenotypes (here, internode allocation fraction and grow time duration).

The structure of our Phytomeric Growth Model is highly generalizable to different species. The Phytomeric Growth Model’s core module, the *Resource Allocation Dynamics* (*RAD),* is a quantization of the temporal coordination of semi-sequential phytomeric growth. Thus, the Phytomeric Growth Model can be adapted to other plants that exhibit phytomeric growth, such as wheat, soybean, and maize. Most models of crop growth treat all leaf biomass as a single pool (e.g., Soybean-BioCro (Matthews et al. 2021), AP-Sim Plant (Brown et al. 2014)). Thus, this Phytomeric Growth Model’s approach could be adapted to better simulate how plant development influences canopy photosynthesis or grain yield.

Although this Phytomeric Growth Model has some similarities to functional-structural plant models (FSPMs), which also simulate explicit phytomers, our approach is simpler, and parameters are easier to estimate using high-throughput phenotypic data in conjunction with computer vision tools, such as PlantCV. Our Phytomeric Growth Model has 15 parameters, with 3 parameters describing allocation fractions. FSPMs are often more complex; e.g. an FSPM for maize has 54 parameters, with 12 parameters describing allocation fractions, and require destructive sampling of aboveground tissue to estimate 12 important parameters (Guo et al. 2006). Although most parameters can be estimated using PlantCV in high-throughput phenotyping systems, preliminary validation is crucial to success when generalizing this Phytomeric Growth Model to identify necessary model structural changes and the appropriate parameter space of that species.

*Caveats.* Developing a model that is mechanistic in structure and developmentally informed, while ensuring phytomeric traits can be estimated with high-throughput phenotypic methods, was challenging. In general, parameter estimation in complex models can be sensitive to initial guesses and parameter boundaries. To address this challenge, we used our initial validation experiment to identify the relevant region of the parameter space, and measurements from our new PlantCV functions to further restrict bounds and obtain reasonable initial parameter guesses, along with global optimization. Even so, translating the highly generalizable Phytomer Growth Model to other species will require additional upfront validation.

As high throughput phenotyping technology continues to expand, including remote sensing, more phenotypic detail and complexity will be detectable (Okada et al. 2024; X. Wang et al. 2019; Tattaris et al. 2016) including traits such as node number and stem length that are features in the Phytomeric Growth Model presented here (Okada et al. 2024). Neural networks, A.I., and deep learning are exciting new approaches to handling the complexity of both plant development and phenotypic image analysis (Zdrazil et al. 2025; Gill et al. 2022). However, such approaches require large amounts of training data, extensive processing, and produce a high number of parameters, making interpretability a challenge (R. Wang et al. 2020). Methods like our Phytomeric Growth Model presented here advantageously couple the development of plant growth in a mechanistic way (R. Wang et al. 2020; Gilpin et al. 2020). Mechanistic models capture feedback mechanisms and emergent system behavior, providing additional biological insight and constraints that are otherwise challenging to express. In the future we anticipate the integration of high-power computational and A.I. approaches with biologically and mathematically appropriate mechanistic models, the coupling of information integration with evolutionarily-favored emergent behaviors (Gilpin et al. 2020).

## Acknowledgements

We thank Andrew D.L. Leakey for his valuable feedback during model development and manuscript preparation.

## Methods

### Validation experiment

#### Experimental Approach

The domesticated *S. italica* (accession B100) was studied alongside its wild relative *S. viridis* (accession A10) in a greenhouse at the Donald Danforth Plant Science Center. This greenhouse was set to a 14 hour day length, with a 28°C day and 22°C night temperature. Relative humidity was maintained between 40-50%. 15 plants of each accession were grown in 1 gal pots. The plants were manually measured, as described below, 3 times a week from 7 days after planting (DAP) until 28 DAP, and then twice a week until dissection, when at least half of the seeds on every plant within each genotype were mature.

#### Manual measurements of phenotypic traits

Near daily measurements of leaf emergence (defined as a visible leaf tip within the whorl) and total leaf number were taken to estimate the phytomer emergence rate. To determine grow time of leaves and internodes, the lengths of leaves and sheaths were measured using a ruler. Sheath length was measured as the distance from one leaf collar to the next leaf collar, or from the soil to the first leaf collar. Sheath elongation was considered to be representative of internode elongation. Leaf length was measured as the distance from the leaf collar to the leaf tip. Each measurement was rounded to the nearest 0.25cm. Once a sheath and leaf measurement remained constant within +/- 0.5cm across three measurement days they were considered fully expanded and were no longer measured. Once emerged, the peduncle length was also measured, from the collar of the last leaf to the bottom edge of the panicle. Panicle length was measured. While *S. italica* has little to no tillering, *S. viridis* grows many tillers, and only the culm was tracked throughout the experiment. For the second experiment, the emergence rate was estimated via visual inspection of Bellwether imaging to identify the emergence of new leaf tips, in addition to final leaf counts in the greenhouse prior to dissection.

#### High-Throughput Imaging Validation Experiment

An experiment used to validate new PlantCV methods was performed. B100 and A10 were grown along with accessions TB_12_0039, TB_12_0265, RIL-13, RIL-68, and RIL-183. The RIL are a recombinant inbred line derived from the cross of B100 and A10. Each line had 10 replicates except for S. italica and S. viridis which each had 5 replicates. These plants were grown in ¼ gal pots, first in a growth chamber, then moved into the Bellwether Foundation Phenotyping Facility (Bellwether Facility), and then into the same greenhouse as the previous experiment. From planting until 12 DAP, plants were in the growth chamber which had a 12 hour day length, day temperatures of 31oC, night temperatures 21oC, with light intensity of 400 (µmol/m2/s) and relative humidity maintained near 40%. The plants were then loaded into the Bellwether Facility. The Bellwether Facility allows for high-throughput phenotyping utilizing a controlled growth chamber environment with a conveyor belt system that moves plants to automatic weighing stations and through an imaging system. Every plant was weighed twice a day and watered to 85% of soil capacity. Imaging occurred once a day using both RGB visible imaging. Plants were photographed from above and at two side angles. The controlled environment was set to nearly the same parameters as the pre-loading growth chamber, with the only difference being the light intensity, set to 500 (µmol/m2/s). Relative humidity also varied more in the facility due to plants moving between the controlled environment and the imaging station, sometimes raising relative humidity levels to above 60%. Plants were grown in the Bellwether Facility for 3 weeks before being moved into the same greenhouse as the previous experiment, where they were grown until seed maturity.

#### Dissections

In all experiments plants were dissected after at least 50% of seeds were mature on all plants within a genotype. Plants were dissected manually using garden shears and razor blades. Leaves and sheaths were removed carefully by hand. Once the leaves and sheaths were removed, internodes were dissected and measured. A cut with shears was made at each node, which was clearly visible, to separate internodes from each other.

### Implementation of the mathematical Phytomeric Growth Model

The mathematical model was implemented in Matlab 2021a. The Resource Allocation Dynamics (*RAD*) module of the Phytomeric Growth Model implements a series of matrix models to describe the timing and allocation of resources to phytomers and phytomer components over time. The Water Usage (*WU*) model uses a system of ODEs to calculate the amount of biomass produced from water provided. See details on model implementation and equations in Supplemental Text 1.

### Implementation of new PlantCV functions

#### Implementation

PlantCV (Gehan et al. 2017) was used to analyze data from (Feldman et al. 2018). Branch point locations were used to assign leaf identity labels per trait extracted, and were tracked to obtain phytomer emergence estimates. Segment lengths were used to track the number of leaves over time. Implementation details in Supplemental Text 2. The resulting estimates were processed as described in Supplemental Text 3. Unrealistic values were removed (e.g., leaf elongation duration <=1, phytomer emergence rate <=1 day). Estimates were processed for QTL analysis in the same method as described below.

### Feldman Data + QTL analysis

Imaging data from (Feldman et al. 2018) was analyzed using PlantCV to extract novel parameter trait measurements as described previously and in Supplemental File 2. Prior to QTL analysis, parameter traits were checked for normality using the Shapiro-Wilks test in R 4.5.1. The normality of each trait transformed by boxcox (from R library MASS) or logarithmic transform were compared, and the transformation that resulted in the highest normality p-value was used for that trait by treatment combination. The genotypic average was used for QTL analysis, performed using Feldman’s foxyQTL pipeline using a non-parametric model (https://github.com/maxjfeldman/foxy_qtl_pipeline).

### Parameter trait estimation

#### Estimating Parameters

Global optimization was implemented in Matlab 2023a using the Multi-Start solver. We solved a set of 200 start points across the parameter space, as defined by the parameter bounds, for each of the 1112 plant samples. 26 samples were removed for missing or incomplete data (less than half of the experimental days had observations of biomass, or the final biomass was less than 3 grams).The Phytomeric Growth Model was fit to the biomass and height data derived from the images, and the water loss data. Water provided data was given as an input to the *WU* module of the model. Parameter bounds were set to be within 15% of the estimated parameter (e.g., phytomer emergence rate, leaf elongation duration, tiller initiation time, and tillering rate). If no PlantCV derived model parameter estimate was available, due to method failure, an average value was used as the initial guess. Randomization settings were retained for reproducibility. Each run results in a function value and parameter set. The water uptake rate frequently reaches lower or upper limit set during parameter estimation, of 1g to 20 g per 0.01 days, or 100g to 2000g of water per day. The water uptake rate is often estimated to be the upper limit during well-watered conditions, and the lower limit during drought conditions. This likely indicates the lack of smaller time-scale information required to fully identify this parameter.

#### Parameter trait analysis

The ability of parameter traits to reflect the underlying biology was additionally assessed using machine learning analyses. Plant size, as measured by final pixel area, was used as the response variable for classification and regression tree analysis (CART) in R (version 4.5.1) using rpart (ver 4.1.24) and partykit (ver 1.2.24). Variable importance was determined using random forest (randomForest package, ver 4.7.1.2) on 500 trees using out-of-bag cross-validation. Both analyses were fit using the data randomly split into training and testing sets, and after verifying fit quality, were run on the full dataset for visualization. Analysis code is provided on the github.

## Supplemental Data

**Supplemental Text 1: Phytomeric Growth Model description** https://drive.google.com/file/d/1uN7nCuqLcPmiGh_DLrg1utBCvlWR24EL/view?usp=sharing

**Supplemental Figures:** phytomer emergence rate (validation experiment), grow time duration (validation experiment), example fits (validation experiment), plantCV vs Phytomeric Growth Model based estimates of the four parameters, banner plot of QTL positions, parameter distributions x treatment condition https://docs.google.com/document/d/1Pqqq15UkjCUncUriETJ7csOYUcT8dspkZJUY8u9lo4s/edit?tab=t.0

**Supplemental Tables:** full QTL list, Phytomeric Growth Model and plantCV; description of tillering, water use parameters by treatment condition (model-based estimates) https://docs.google.com/document/d/1SoZlBQqIYHxLZQZXhqstGdaF41kjU8EFBaZug8OJg44/edit?tab=t.0

