## Supplemental Tables for "Modeling phytomeric growth using high-throughput phenotyping deconvolutes genetic control of complex traits underlying size and water use in *Setaria*"

| Compartment | Parameter Trait (units) | Value |
| --- | --- | --- |
| **Water Use (WU)** | Water uptake (g / day) | Estimated |
| | Marginal WUE (g biomass produced / g leaf) / (g water available * day) | Estimated |
| **Relative Allocation Fractions (RAF)** | Internode allocation | Estimated |
| | Sheath allocation | *Held equal to internode alloc.* |
| | Reproductive allocation | Estimated |
| | Peduncle allocation | Fixed, 0.01 |
| | Initial Size Multiplier | Estimated |
| **Resource Allocation Dynamics (RAD)** | Phytomer emergence rate (per day) | Estimated† |
| | Leaf elongation duration | Estimated† |
| | Internode elongation duration | Estimated |
| | Sheath elongation duration | *Held equal to internode e.d.* |
| | Reproductive elongation duration | Fixed at 1.5 times leaf elongation duration* |
| | Panicle emergence time (DAP) | Estimated |
| **Tillering Deconvolution (TD)** | Tiller start (DAP) | Estimated† |
| | Tiller rate ($\beta_1$) | Estimated† |
| | Tiller rate intercept ($\beta_0$) | Fixed to PlantCV-estimated value† |

**Table 1. Parameter traits.** All values are estimated on a per-sample basis, except where noted. Parameters are unitless except where noted. See main text for discussion of the marginal WUE. † indicates value is estimated by PlantCV. * *Very few plants reach full elongation of the peduncle during this experimental timecourse, and this value is largely unused*.

| Compartment | Trait | PlantCV QTL | Model QTL |
|---|---|---|---|
| **Water Use (WU)** | Water uptake (g / 0.01 day) | – | 5@110*<br>9@35* |
| | Marginal WUE (g biomass / g leaf) / (g water * time) | – | 2@95*<br>3@47*<br>5@101**<br>7@100*<br>9@34* |
| **Relative Allocation Fractions (RAF)** | Internode allocation | – | 2@87<br>7@28<br>7@101<br>8@41* |
| | Reproductive allocation | – | 8@6<br>5@100* |
| **Resource Allocation Dynamics (RAD)** | Phytomer emergence rate (per day) | N/A | |
| | Leaf elongation duration | N/A | 2@101 |
| | Internode elongation duration | – | 5@81* |
| | Panicle emergence time (DAP) | – | 8@6 |
| **Tillering Deconvolution (TD)** | Tiller start (DAP) | 2@95*<br><br>7@100* | 2@95*<br>5@59*<br>7@100*<br>9@39* |
| | Tiller rate | 1@43**<br>2@9<br>2@111*<br>3@5**<br><br>5@85*<br><br><br>6@74*<br>7@51* | 1@43**<br>2@9**<br><br>3@5**<br>4@81<br><br>5@100*<br>6@48<br>6@64*<br>7@42** |
| **Calculated Values** | Tiller fraction of the total biomass | – | 5@104*<br>7@42*<br>7@96*<br>9@34* |
| | Number of leaves | – | |

| | | | |
|---|---|---|---|
| | Number of Tillers | – | 3@5*<br>5@105*<br>6@73<br>7@51* |

**Table 2. Full QTLs for parameter traits.** The symbol * indicates wet conditions, and the symbol ** indicates QTL found for both wet and dry conditions. Parameters are unitless except where noted.

| | **Dry** | **Wet** |
|---|---|---|
| **Tiller start (DAP)** | 21 +/- 6.6 days | 16.7 +/- 3.9 days |
| **Tiller rate** | 0.99 +/- 0.3 | 1.64 +/- 0.4 days |
| **Tiller fraction of total biomass** | 0.48 +/- 0.5 | 0.82 +/- 0.5 |
| **Number of Tillers** | 1.1 +/- 0.6 | 2.1 +/- 1.1 |
| **Marginal WUE** | 0.0015 +/- 0.0006 | 0.0009 +/- 0.0003 |
| **Water uptake rate** | 15.4 +/- 7.5 | 11.3 +/- 6.5 |
| **Phytomer emergence rate** | 1.9 +/- 0.1 | 1.9 +/- 0.7 |

**Table 3. Descriptions of model parameter estimates.** We found that tiller start time was 17 days after planting on average in wet conditions, and 21 days after planting during drought conditions. Tiller start time, tillering rate, and the tiller fraction of total mass all significantly decreased in drought conditions across genotypes ($p < .0001$, paired t-test, unequal sample variance). No correction was done due to the very low p-values.

Marginal WUE and the water uptake rate were both significantly different across treatment conditions ($p < .0001$ via t-test). Water uptake rate was estimated to be 11.3 grams per 0.01 days (ie, 15 mins) in well-watered conditions, and 15.4 grams per 0.01 days in drought conditions. The phytomer emergence rate was also significantly different across treatment conditions ($p < .0001$).

| Dependent Variable | *t* | *df* | *p* | *d* | 95% CI |
|---|---|---|---|---|---|
| intBeta | 2.64 | 1,087.27 | .008** | 0.16 | [0.04, 0.28] |
| Rho | -0.16 | 1,104.79 | .876 | -0.01 | [-0.13, 0.11] |
| toInt | -5.37 | 793.89 | < .001*** | -0.32 | [-0.44, -0.20] |
| leafBeta | -2.45 | 837.04 | .015* | -0.15 | [-0.26, -0.03] |
| waterUptakeRate | -4.01 | 1,004.23 | < .001*** | -0.24 | [-0.36, -0.12] |
| marginalWUE | 27.01 | 855.22 | < .001*** | 1.62 | [1.49, 1.76] |
| panEmerg | -1.84 | 1,109.20 | .066 | -0.11 | [-0.23, 0.01] |
| PhytEmergenceRate | 0.72 | 897.43 | .471 | 0.04 | [-0.07, 0.16] |
| tillerStart | 13.24 | 831.24 | < .001*** | 0.79 | [0.67, 0.92] |
| tillerRate | -32.10 | 905.63 | < .001*** | -1.92 | [-2.07, -1.78] |
| numLeaves | -1.60 | 1,107.09 | .110 | -0.10 | [-0.21, 0.02] |
| tillerFractionTotalMass | -21.30 | 1,074.70 | < .001*** | -1.28 | [-1.41, -1.15] |
| numTillers | -17.71 | 718.64 | < .001*** | -1.06 | [-1.19, -0.93] |

**Supplemental Table 4. T-Test results comparing parameter estimates across genotypes in well-watered and drought conditions.** Almost all parameters show a significant change under drought conditions. Large effect sizes (Cohen's *d* > 0.8) are seen in tiller start time, tillering rate, tiller fraction of total mass, the marginal WUE, and the number of tillers. The value of *t* indicates the direction of change going from well-watered to drought conditions.

| | mWUE | tillerRate | Total water | bWUE | totalPixels |
|---|---|---|---|---|---|
| WW, dry weight | **0.68** | 0.45 | **0.81** | 0.19 | **0.61** |
| WD, dry weight | **0.57** | 0.25 | **0.68** | 0.25 | **0.51** |
| WW, totalPixels | **0.60** | **0.54** | **0.64** | -* | - |
| WD, totalPixels | 0.39 | 0.39 | 0.43 | -* | - |

**Supplemental Table 5. Model-based estimates of marginal WUE have comparable correlation coefficients to biomass estimates compared to dry weight measurements.** Moderate (0.5 - 0.7) to high correlation coefficients (0.7+) are bolded. * totalPixels and totalWater are used to calculate the imaging-based estimate of the whole-plant WUE ("bWUE", totalPixels/totalWater), which naturally produces uninteresting, strong correlations in the estimates. Marginal WUE is as strong or more strongly correlated to the dry weight measurements compared to whole-plant WUE ("bWUE") and totalPixels, the imaging-based estimates of plant biomass.
