## Supplemental Figures for "Modeling phytomeric growth using high-throughput phenotyping deconvolutes genetic control of complex traits underlying size and water use in *Setaria*"

### Supplemental Figures: Greenhouse Validation Experiment

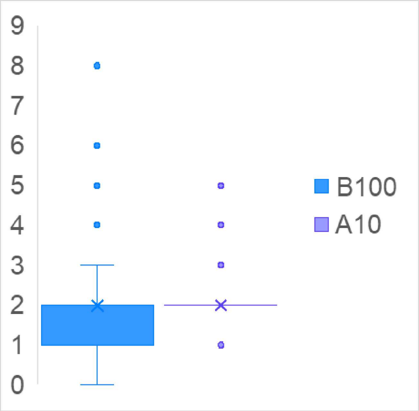


**Supplemental Figure 1. Emergence rates for the two genotypes in the manual validation experiment.** The phytomer emergence rate was tracked by hand every 2-3 days until panicles emerged for all plants.

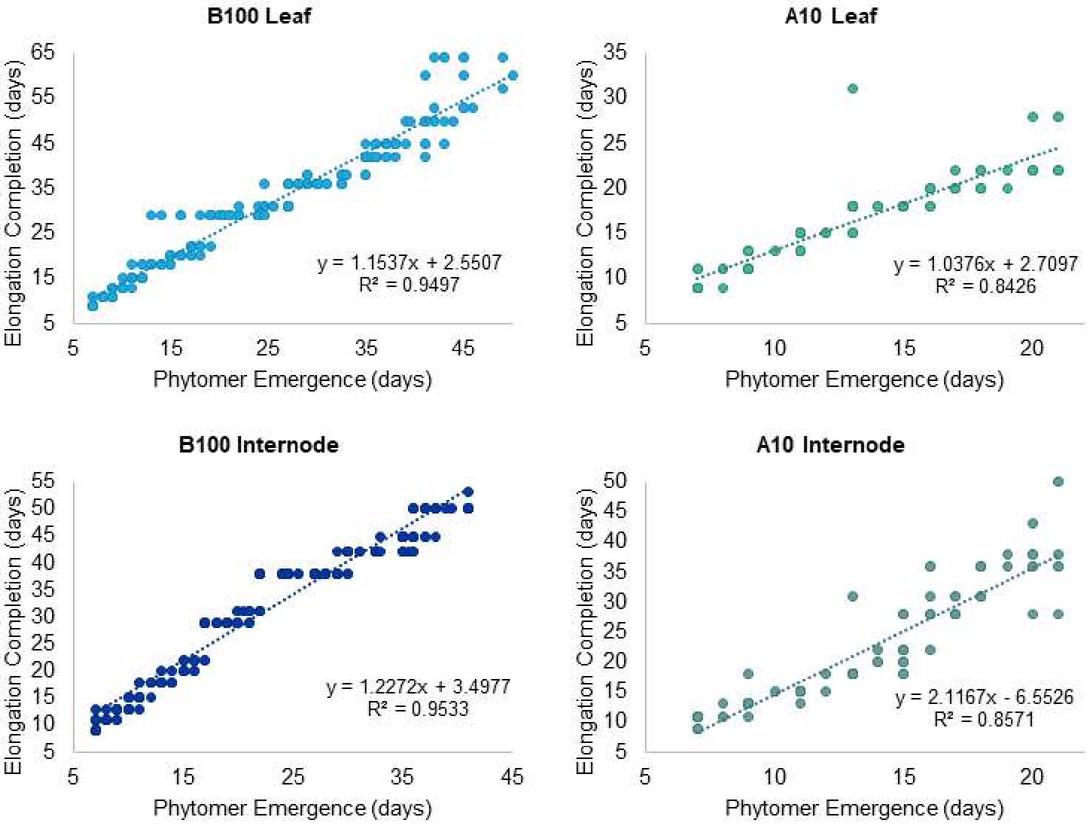


**Supplemental Figure 2. Linear regressions describe the relationship between emergence date and elongation completion date to determine the growth multiplier parameter.** Linear regressions were applied to the phytomer emergence and elongation completion dates, pooled by genotype (A10 or B100) across phytomer component (leaf or internode).

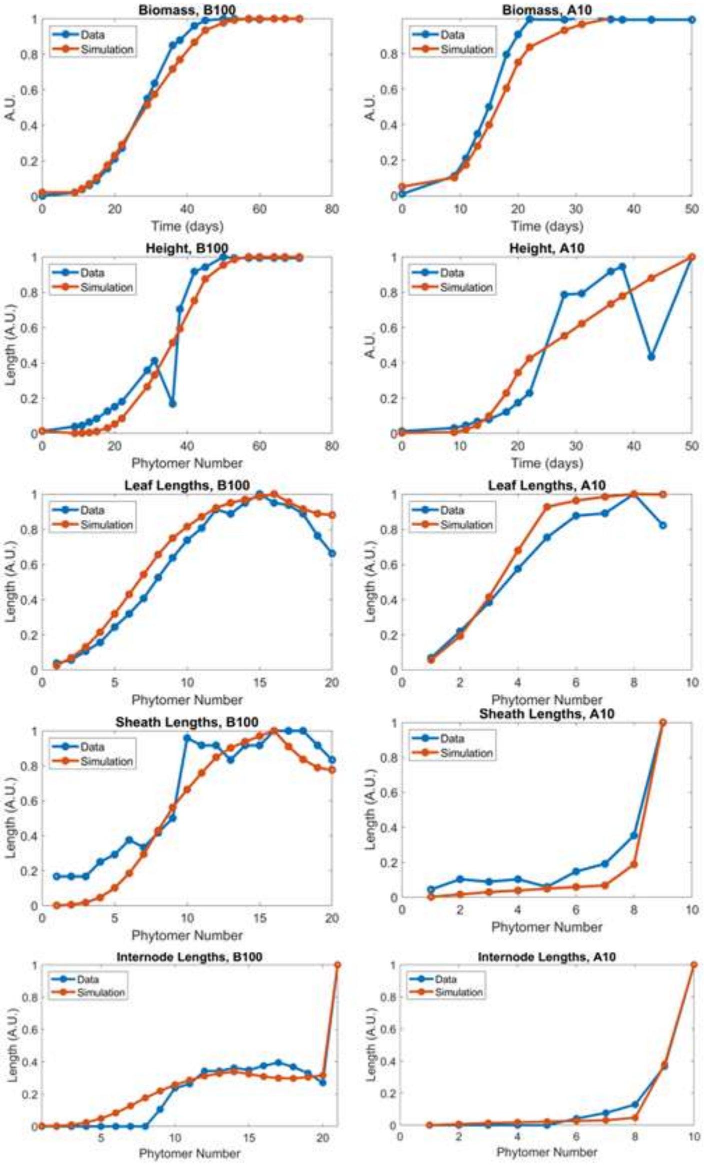


**Supplemental Figure 3. Example model fits to the manual validation experiment.** Data from a single randomly chosen B100 plant shown (blue) with model fit (orange). A) Biomass growth over time, approximated using the sum of leaf lengths. B) Height over time, approximated using the sum of sheath lengths plus the peduncle. C) Leaf blade lengths, normalized. D) Leaf sheath lengths, normalized. E) Internode and peduncle lengths, normalized.

### Supplemental Figures: High-Throughput Experiments

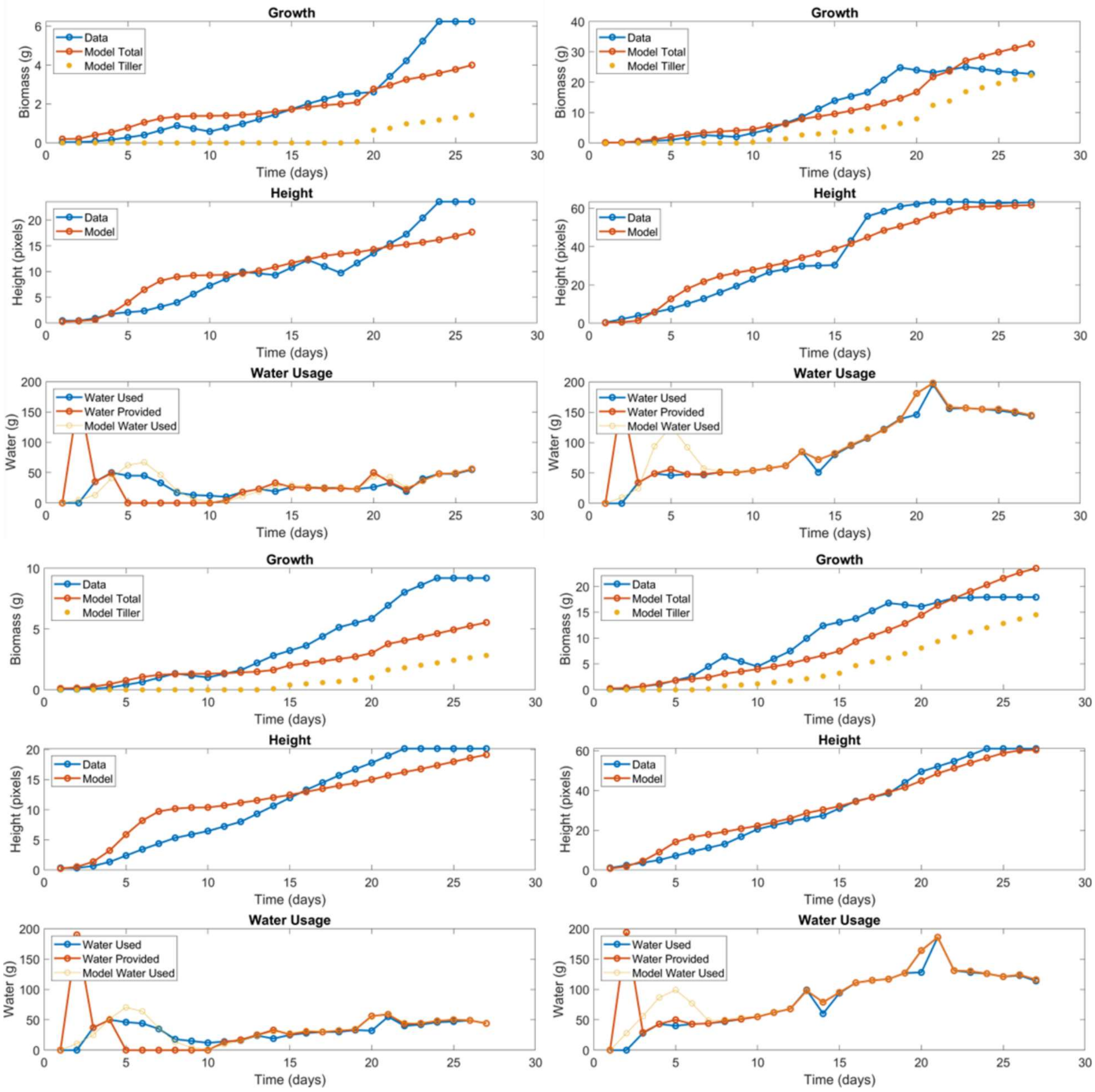


**Supplemental Fig 4. Examples of model performance on high-throughput phenotyping data.** Example fits in drought (left column) and well-watered (right column) conditions. A random 10% of samples were plotted after parameter estimation, and the first two of each treatment condition are shown here. Clockwise from top left, the genotypes presented here are A10, RIL11, RIL33, and RIL16. The model breaks down total total biomass (measured as pixels in PlantCV, and converted to grams using data obtained in Feldman et al 2018) into culm and tiller biomass. Water provided data is passed to the model, and the model calculated water usage per day is fit to the water used data. Additional figures can be found on the Phenomodel github.

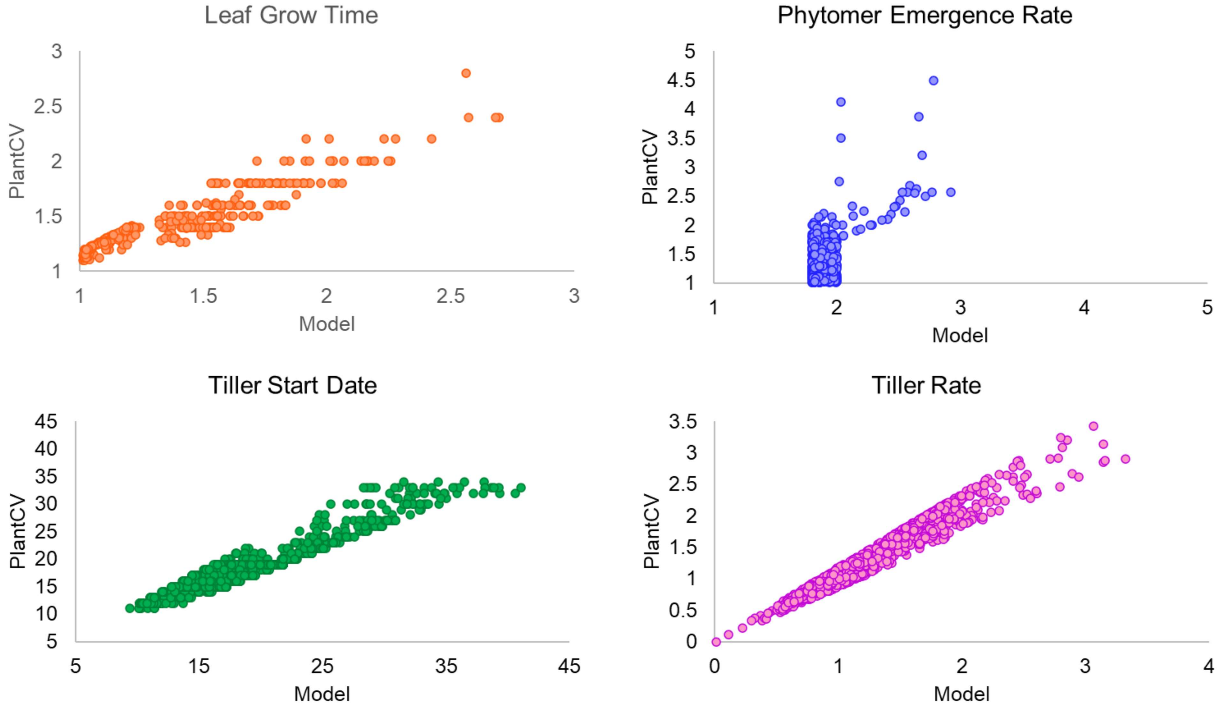


**Supplemental Figure 5. How the model modified initial plantCV estimates.** Leaf elongation time was adjustable within 15% of plantCV estimate. If leaf elongation time was not able to be estimated by plantCV, it was set to 1.2, the approximate value observed in the validation experiment. Leaf elongation duration had a lower limit of 1.01. For the phytomer emergence rate, unrealistic values were filtered and adjusted, and values within 15% of the estimate, with the estimate constrained to be within 1.5 and 3 days. PlantCV phytomer emergence rate estimates were frequently unrealistic given the presence of a panicle, for example, providing a time constraint of all phytomers emerging within the 30 days of the experimental time course. If a panicle was observed, phytomer emergence rate received an upper limit of 2.5 days. Tiller initiation time was within 15% of the plantCV estimate. If tillering initiation time was estimated to be earlier than or equal to 2 days after the start of the experiment, it was adjusted to 3 days after the start of the experiment to account for possible mis-categorization due to small plant size in plantCV. If tiller initiation time was equal to the end of the experiment (i.e., no tillering observed), then the bounds were set to range from 5 days prior to the end of the experiment to the end of the experiment + 9 days to allow for potential missing data. Tillering rate was within 15% of the estimate. If tillering rate was not able to be estimated, then tiller initiation time was bounded between 0.1 and 3.

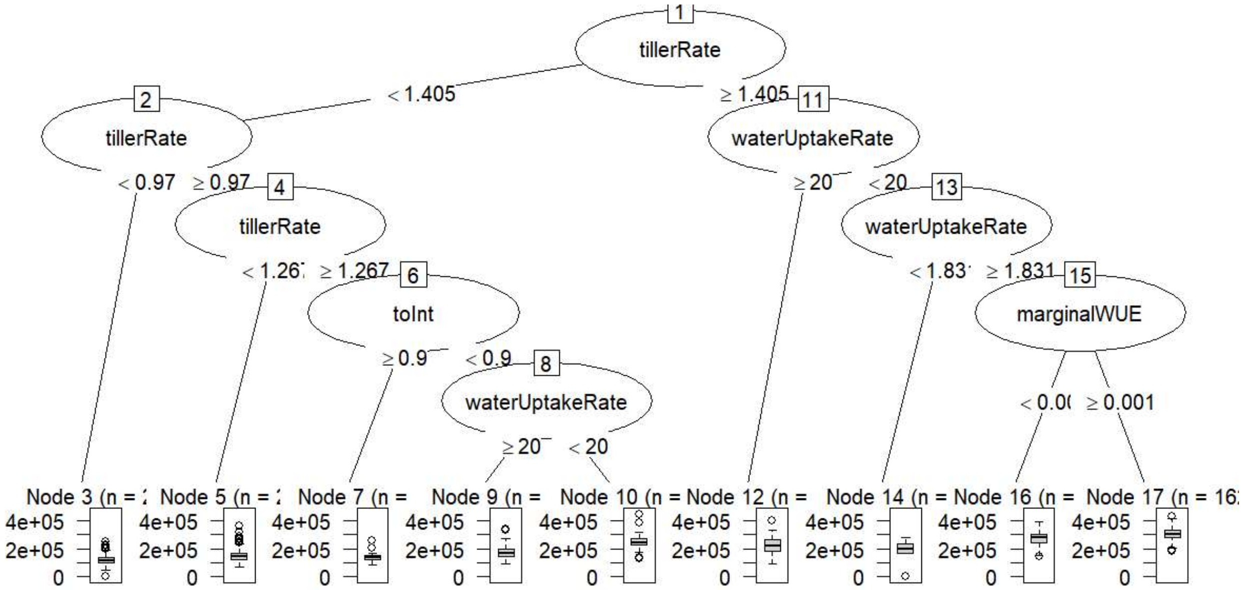


**Supplemental Figure 6. Plant size across both treatment conditions can be characterized using tillering and water usage parameters.** Machine learning based clustering (classification and regression tree, CART) suggests that the tillering rate and marginal WUE are highly predictive of plant size across both treatments. Larger plants (as measured by total pixels, the boxplots shown in the bottom row of the tree) tend to have higher tillering rates (≥1.405) and higher marginal WUE (≥ 0.001). Both treatment conditions are represented here. Only model parameters and calculated values (e.g., number of leaves, tiller fraction of total mass) are included in this model.

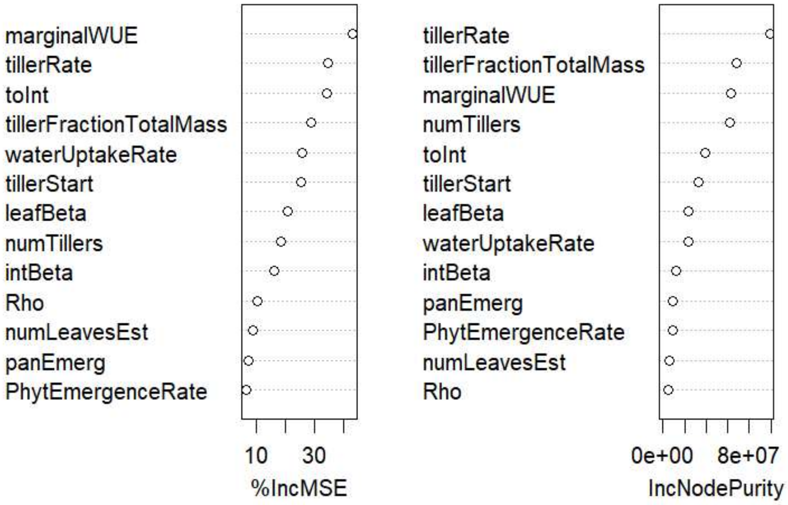


**Supplemental Figure 7. The marginal water use efficiency and the tillering rate are influential parameters driving plant responses.** A random forest model was used to analyze the influence of model parameters on plant size across both treatment conditions and all genotypes. The marginal water use efficiency and the tillering rate, as well as the allocation to the internodes and the tiller fraction of total mass, are identified as highly influential parameters.
