## Supplemental Text for "Modeling phytomeric growth using high-throughput phenotyping deconvolutes genetic control of complex traits underlying size and water use in *Setaria*"

### 1 Model Description

The model is based on two core hypotheses. The first is that the growth and development of each phytomer is determinate and that it ends after a fixed duration, presumably set by the plant's genetics. The second is that the resources needed for growth are only allocated to growing phytomers and each growing phytomer receives an equal fraction of the total resource supply. The ultimate size of a given phytomer depends on the total resource supply during its development and the number of other phytomers that were growing concurrently.

In this model, each phytomer consists of a leaf and stem internode. The leaf is split into two components: its lamina (the flat part) and its sheath (i.e., *Setaria*) or petiole (i.e., *Glycine*). Thus, phytomers have three component tissues (lamina, sheath/petiole, and internode). Each component develops at a different rate, but the allocation of resources follows the core hypotheses. As *Setaria* produces tillers, each tiller adds another set of phytomers whose development is modeled in the same way as the main shoot. Finally, the model includes reproductive structures (e.g., the panicle in *Setaria*, the pods in *Glycine*) as separate "phytomers". In all cases, allocation is determined by the number of actively growing phytomer components. As the model was first developed for Setaria, we use Setarias anatomy when describing the model.

#### 1.1 Growth and Development

To describe the growth and development of phytomers, let $t$ denote the number of days after planting (DAP). Let $t_g$ be the germination date, defined as the DAP at which the first phytomer of the main shoot has emerged. In the greenhouse experiment, measurements began at $t = 7$ DAP and after the emergence of the first phytomer, so we estimate $t_g \approx 5$ DAP. A new phytomer of the main shoot (the culm) emerges every $E$ days, so the second phytomer emerges at age $t_g + E$, the third at age $t_g + 2E$, and so on. Thus, the age of emergence $e_j$ of the $j$-th phytomer of the main shoot occurs $e_j = t_g + (j-1)E$. The panicle of the main shoot emerges $E$ days after the last phytomer, so if the main shoot has $n$ phytomers in total, then the panicle's age of emergence is $e_{\text{panicle}} = t_g + N_{\text{phyt}}E$ DAP. *Setaria* produces tillers, so the phytomers of a tiller's shoot follow the rules for development. Tillering is assumed to begin at a certain age $t_1$ and a new tiller is produced every $T$ days. The $i$-th tiller's shoot emerges at a age $t_i = t_1 + (i-1)T$, the date at which its first phytomer has emerged, then produces a new phytomer every $E$ days.

Each component tissue of each phytomer (i.e., the lamina, sheath, and internode) grows a set duration of time, called its development time $D$. The $j$-th phytomer begins growing at its age of emergence $e_j$ DAP, its lamina finishes growing at $e_j + D_{j,\text{lamina}}$ DAP, its sheath at $e_j + D_{j,\text{sheath}}$ DAP, and so on. In the *Setaria* model, the development time is proportional to the plant's age, so that it increases with age and thus younger phytomers have longer development times than older phytomers. Namely, the development time $D_{j,k}$ of the $j$-th phytomer's component $k$ is $e_j + D_{j,k} = \beta_k e_j$. The development time is determined by a component-specific parameter $\beta_k$, the development time multiplier. The parameter $\beta_k$ is constant in time and the same for all phytomers of all shoots (the culm and any tillers).

Thus, the model encapsulates the development of the plant into a set of parameters which define the rate of shoot and phytomer development: the emergence rate $E$, the development time multiplier $\beta_k$, the age of first tiller $t_1$, the tillering rate $T$, and the total number of phytomers per shoot before reproduction. These development parameters determine when phytomers are growing, but they do not entirely determine the size of the phytomers. The ultimate size of each phytomer depends on resource supply during.

| Parameter | Unit | Definition |
|---|---|---|
| Emergence rate $E$ | days | A new phytomer is produced every $E$ days. |
| Development time multiplier $\beta_k$ | – | The ratio of a given phytomer's component $k$ age of maturity to its age of emergence $e_j + D_{j,k}/e_j = \beta_k$ |
| Tillering rate $T$ | days | A shoot for a tiller is produced every $T$ days |
| Age of first tiller $t_1$ | DAP | The first phytomer of the first tiller emerges $t_1$ DAP |
| $N_{\text{phyt}}$ | – | The number of phytomers on a shoot before its panicle emerges |

#### 1.2 Resource Allocation Dynamics

The model defines resource allocation dynamics as follows. Suppose the plant has a supply $R$ of some resource. For clarity of exposition, $R$ is essentially the rate of biomass production from photosynthesis. Later, we discuss modifications necessary for the high-throughput experiments. Let $m_{jk}(t)$ be the fresh biomass of component $k$ of phytomer $j$ of any shoot, and let $w_{jk}$ be the fraction of biomass production $R$ allocated to component $k$ of phytomer $j$ so that:

$$\frac{dm_{jk}}{dt} = w_{jk}(t)R(t) \tag{1}$$

The model's core hypothesis is that the growing components of each phytomer receive the same fraction of $R$ that each developing phytomer receives an equal fraction of the total resource so that the resource allocation changes dynamically as the number of active-developing phytomers changes. However, allocation is not equal between the components.

The formula for $w_{jk}$ depends on the number of developing or growing components at time $t$. To count them, we define a growth indicator function $g_{jk}(t)$ such that $g_{jk}(t) = 1$ if component $k$ of phytomer $j$ is growing at time $t$, and $g_{jk}(t) = 0$ if not. This indicator function is determined by the growth and development parameters (described above), independent of resource availability. A concise formula for $g_{jk}$ is given in Equation (2) for the first shoot (the culm). The formula for a tiller's shoot is the same but $t_g$ is replaced with its date of emergence. To simplify the description, we will assume no tillers for this section.

$$g_{jk}(t) = \begin{cases} 1 & u \in [0,1] \\ 0 & u \notin [0,1] \end{cases} \qquad \text{where} \quad u = \frac{t - (t_g + (j-1)E)}{(\beta_k - 1)\,(t_g + (j-1)E)} \tag{2}$$

Summing $g_{jk}$ over all phytomers gives a count $G_k$of the number of phytomers whose component $k$ is growing at time $t$.

$$\#\text{ of growing component } k = G_k(t) = \sum_j g_{jk}(t)$$

Let $a_k > 0$ be a parameter for each component $k$ (i.e., lamina, sheath, internode, reproductive structure, etc.) called the allocation weight of component $k$ so that the allocation fraction $w_{jk}$ is given by (3).

$$w_{jk}(t) = \frac{a_k g_{jk}(t)}{S(t)} \qquad S(t) = \sum_{jk} a_k g_{jk}(t) \tag{3}$$

This formula completely defines the resource allocation dynamics.

To show how this formula satisfies the core model hypotheses, note that the allocation weights $a_k$ control the ratio of allocation between components. Sum over the phytomers to obtain the allocation fraction to component $k$.

$$\text{allocation to component } k = \sum_j w_{jk}(t) = \frac{a_k G_k(t)}{S(t)} \qquad S(t) = \sum_k a_k G_k(t)$$

This allocation fraction is a weighted average of the number phytomers with whose component $k$ is growing at time $t$. Note that because components develop at a different rate, $G_k$ is not necessarily the same for all components. The ratio of the component $k$ allocation fraction to the component $k'$ allocation fraction is:

$$\frac{\text{allocation to component } k}{\text{allocation to component } k'} = \frac{\sum_j w_{jk}(t)}{\sum_j w_{jk'}(t)} = \left(\frac{a_k}{a_{k'}}\right)\left(\frac{G_k(t)}{G_{k'}(t)}\right)$$

So this ratio equals the product of the ratio $a_k/a_{k'}$ and the ratio of the number of developing components. For instance, if $a_{\text{lamina}} = 1.6$ and $a_{\text{sheath}} = 0.8$, then allocation to the lamina will be twice the allocation to the sheath if there is an equal number of developing lamina and sheath, but in general, the number of growing lamina to growing sheaths is not equal to one, as it depends on how quickly they develop relative to each other.

Note this formula satisfies the core hypothesis that all growing phytomers receive an equal fraction of the total resource supply:

$$\text{allocation to component } k \text{ on phytomer } j = \frac{w_{jk}(t)}{\sum_i w_{ik}(t)} = \frac{a_k g_{jk}(t)}{a_k G_k(t)} = \frac{1}{G_k(t)}$$

The last equality only holds if phytomer $j$ has a growing component $k$.

#### 1.3 Estimating biomass production in the validation experiment

As described in the main text, the validation experiment was used to determine the parameters of the growth and development model, and validate the model's ability to describe resource allocation. For this experiment, the biomass production $R(t)$ was estimated from empirical observations as follows.

Given a rate of biomass production $R(t)$, the resource allocation dynamics model, equation (**??**), predicts the amount of biomass allocated to each phytomer and component. To estimate $R$, the biomass $m_{jk}$ of each phytomer was assumed to be proportional to the measured length $x_{jk}$ of the leaf lamina. Thus, the total biomass is proportional to the total leaf length $x = \sum_{jk} x_{jk}$, and the change in total leaf length is proportional to biomass production $dx/dt \propto dm/dt$. To standardize comparisons between the model and observations from real plants, the observed leaf length of each phytomer was normalized to the fraction of total leaf length produced $\max(x)$. The resource allocation dynamics model defines allocation up to a multiplicative constant, so the normalized has no influence on the model's predictions for allocation. Thus, the accumulated biomass could be estimated from measured leaf length and approximated using a Hill function model:

$$\frac{x(t)}{\max(x)} = \left(1 + \frac{rt_f}{t}\right)^{-H} \qquad t_f = \beta_{\text{lamina}} \left(E(N_{\text{phyt}} - 1) + t_g\right) \tag{4}$$

where $r$ and $H$ are free parameters, the growth rate and cooperativity coefficient respectively, fit to observed growth when data is available. The quantity$t_f$ is the time when all vegetative phytomers of the main shoot have finished growing. The rate of biomass production is then proportional to:

$$\frac{d}{dt}\left(1 + \frac{rt_f}{t}\right)^{-H}$$

In the model, if all vegetative phytomers finish growing before reproductive growth begins, then the final length of the internode, sheath, or leaf of any phytomer does not depend on the reproductive growth. The development time multipler of the reproductive phytomer is set as $1.5\beta_{\text{leaf}}$.

#### 1.4 Estimating biomass production in the high-throughput experiment

The high-throughput phenotyping system records images of each plant through time rather than measurements of biomass; however, the plant's size was estimated as the number of pixels in each image. As Feldman *et al.* (2018) weighed the final biomass of most plants, we could pair a pixel count with final biomass. We used the mean biomass divided by mean pixel count[1] to obtain a conversion factor (gram / pixel). Using this conversion factor, we converted pixel counts from all images into biomass estimates, and these estimates were used in all subsequent calculations.

Rather than estimating the biomass production directly, a system of ordinary differential equations (ODEs) was used to dynamically simulate water use rate and biomass production. Biomass production was modeled as a "trade" of water for biomass. Let $W$ denote the amount of water in the pot's soil. The plant takes up water at a rate $U = uW$, proportional to the amount of water in the soil. The parameter $u$, the water uptake coefficient, determines how quickly water depletes from the soil. The rate of water uptake determines the amount of water available for photosynthesis via leaf transpiration. Let $X$ denote the plant's water balance defined as the cumulative water uptake minus the cumulative transpiration, then $dX/dt$ equals the rate of water uptake minus the rate of transpiration.

Each gram of water transpired yields $\varepsilon$ grams of biomass. We call $\varepsilon$ the water use efficiency (WUE) but note that the term water use efficiency often refers to other related quantities (such as the rate of CO2 assimilation per transpiration). When shading is negligible, the rate of photosynthesis (i.e., $CO_2$ assimilation) increases proportionally to leaf mass, and therefore the rate of transpiration is proportional to leaf mass. The rate of transpiration from the culm is assumed to be $\rho X M_{\text{c,leaf}}$ where $M_{\text{c,leaf}}$ is the total leaf mass of the culm. Similarly, the rate of transpiration from all tillers is $\rho X M_{\text{t,leaf}}$, proportional to leaf. The parameter $\rho$ controls the rate of transpiration and corrects the units. Thus, the rate of biomass production is $dM_{\text{c}}/dt = \varepsilon\left(\rho X\right) M_{\text{c,leaf}}$ for the culm and $dM_{\text{t}}/dt = \varepsilon\left(\rho X\right) M_{\text{t,leaf}}$ for the tillers. The allocation of biomass to each phytomer of the culm then follows the same dynamics in the validation experiment, governed by the growth and development model as described above in equation (1).

[1]Or do you mean the mean (biomass per pixel count)?

$$\frac{dW}{dt} = -uW \tag{5}$$

$$\frac{dX}{dt} = uW - \rho X(M_{\mathrm{c,leaf}} + M_{\mathrm{t,leaf}}) \tag{6}$$

$$\frac{dM_{\mathrm{c}}}{dt} = \varepsilon\,(\rho X)\,M_{\mathrm{c,leaf}} \tag{7}$$

$$\frac{dM_{\mathrm{t}}}{dt} = \varepsilon\,(\rho X)\,M_{\mathrm{t,leaf}} \tag{8}$$
